# Pentose Phosphate Pathway Protects *E. coli* from Antibiotic Lethality

**DOI:** 10.1101/2024.09.07.611820

**Authors:** Tatyana Seregina, Rustem Shakulov, Konstantin Shatalin, Svetlana Sklyarova, Irina Petrushanko, Vladimir Mitkevich, Alexander Makarov, Alexander S. Mironov, Evgeny Nudler

## Abstract

Disruption of both branches of the canonical pentose phosphate pathway (PPP) in *E. coli* by combined inactivation of the *zwf* and *talAB* genes provokes the restoration of the ancient anabolic variant of PPP (aPPP). In the aPPP, pentose-5-phosphates are synthesized unidirectionally from fructose-6-phosphate and glyceraldehyde-3-phosphate by transketolase B, aldolase A, and phosphatase GlpX, converting sedoheptulose-1,7-bisphosphate to sedoheptulose-7-phosphate. Unexpectedly, the double *zwf talAB* mutant exhibits decreased survival after treatment by diverse classes of antibiotics with little effect on the minimal inhibitory concentration. Simultaneously, we found that killing effect of antimicrobials on the *zwf talAB* mutant could be reversed by the inactivation of either *purR* or *deoB* genes, both responsible for ribose-5-phosphate content in the mutant strain. Enhanced biosynthesis of the cell wall component ADP-heptose from sedoheptulose-7-phosphate also suppressed killing effect of antibiotics on the *zwf talAB* mutant. Furthermore, the inactivation of the Entner-Doudoroff pathway (*Δedd*) or shifting the metabolic equilibrium by the addition of exogenous phosphogluconate reverts aPPP to glycolysis, preventing the accumulation of excess pentose phosphates and the occurrence of the futile cycle in *zwf talAB* cells, thus desensitizing them to antibiotics. Our findings show that ribose-5-phosphate metabolism plays a crucial role in bacterial tolerance to a wide range of bactericidal antibiotics. We propose that targeting PPP could be a promising strategy for developing new therapeutic agents aimed at potentiating clinically significant antimicrobials.

**IMPORTANCE:** Recent studies have revealed the crucial role of bacterial cell’s metabolic status in its susceptibility to the lethal action of antibacterial drugs. However, there is still no clear understanding of which key metabolic nodes are optimal targets to improve the effectiveness of bacterial infection treatment. Our study establishes that the disruption of the canonical pentose phosphate pathway induces one-way anabolic synthesis of pentose phosphates (aPPP) in *E. coli* cells, significantly increasing the killing efficiency of various antibiotics. It is also demonstrated that the activation of ribose-5-phosphate utilization processes restores bacterial tolerance to antibiotics. We consider the synthesis of ribose-5-phosphate to be one of the determining factors of bacterial cell stress resistance. Understanding bacterial metabolic pathways, particularly the aPPP’s role in antibiotic sensitivity, offers insights for developing novel adjuvant therapeutic strategies to enhance antibiotic potency.

## INTRODUCTION

It is now recognized that the ability of antibiotics to eradicate bacteria is closely tied to their metabolic state, encompassing biosynthetic rates, respiration, and redox balance (1–5). Nonetheless, the specific metabolic mechanisms influencing antibiotic tolerance remain incompletely understood.

The pentose phosphate pathway (PPP) serves as a pivotal metabolic hub for pentose phosphate synthesis. The canonical PPP comprises two branches: oxidative decarboxylation of glucose-derived phosphogluconate, generating NADPH for anabolic processes and oxidative stress defense; and the non-oxidative branch, which lacks a driving force-generating reaction for pentose phosphate production (6). Certain organisms, such as *Prevotella copri*, utilize modified non-oxidative PPP variants to assimilate one-carbon sources like xylose (7–13). This includes anabolic aPPP, where fructose-6-phosphate and glyceraldehyde-3-phosphate are converted to sedoheptulose-1,7-bisphosphate and sedoheptulose-7-phosphate using aldolase A and phosphatase GlpX enzymes, bypassing transaldolase (Fig. 1) (7, 14).

**Fig. 1.**
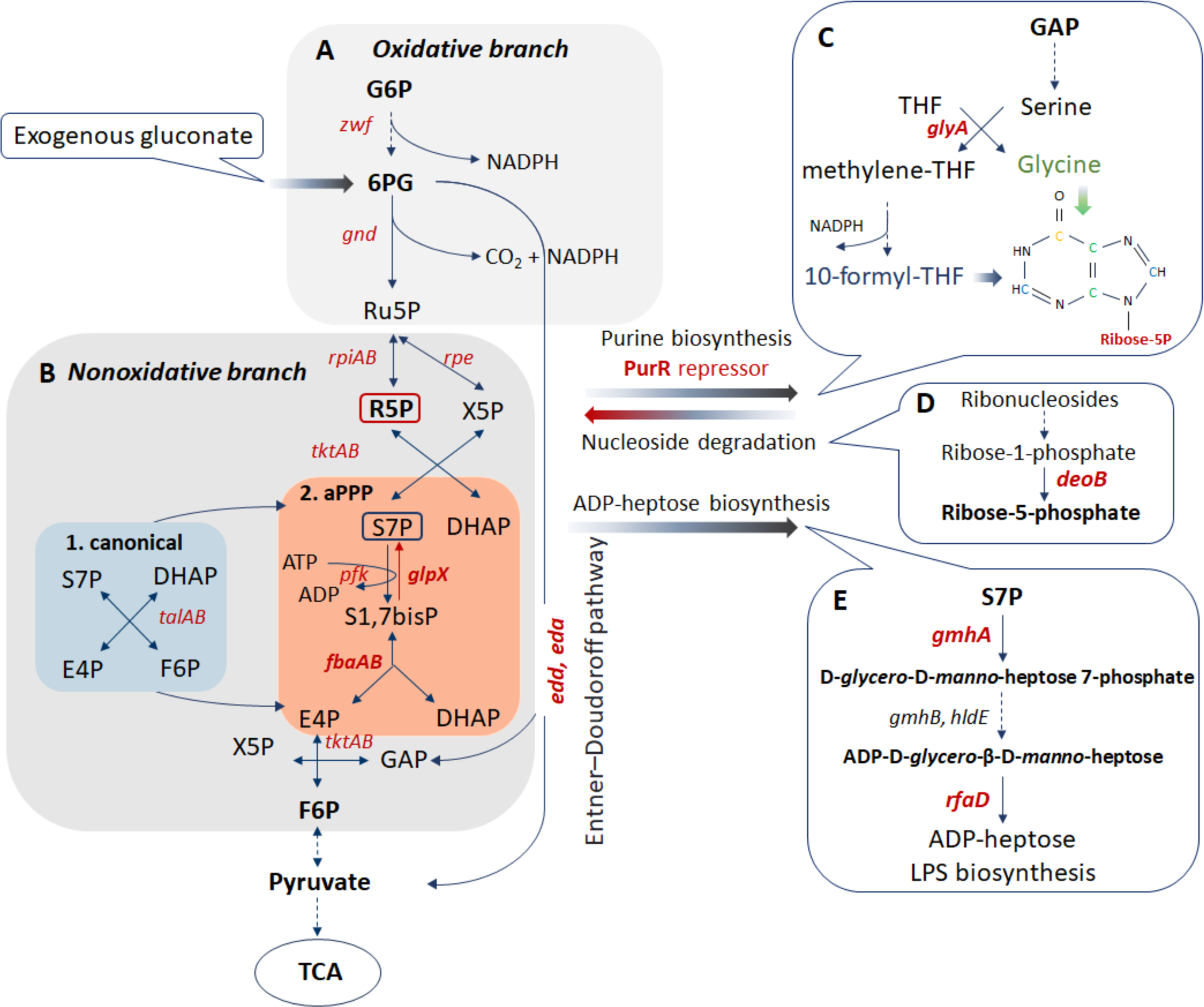
Scheme demonstrating the processes of synthesis and assimilation of ribose-5-phosphate in *E. coli* cells. The anabolic synthesis of pentose phosphates (aPPP) occurs in *E. coli* upon inactivation of the oxidative branch by *zwf* mutation (A) and replacement activity transaldolase TalAB (B1) by the glycolysis enzymes aldolase FbaA, phosphatase GlpX, and phosphofructokinase Pfk (B2). The futile cycling of S1,7P and S7P formed in aPPP is indicated by an orange square. (C) Derepression of purine nucleotide synthesis by inactivation of the PurR repressor enhances the efflux of ribose-5phosphate from the PPP. The serine-glycine pathway is also under the control of PurR and is another source of generation of reducing equivalents of NADPH. (D) Catabolism of ribonucleosides makes a significant contribution to the intracellular pool of ribose-5-phosphate. The final stage of the conversion of ribose-1-phosphate to ribose-5-phosphate is carried out by phosphopentomutase *deoB*. (E) Sedoheptulose-7-phosphate, which is involved in the formation of the futile cycle of aPPP, serves as a precursor to the lipopolysaride component ADP-heptose. Abbreviations used: G6P – glucose-6-phosphate, 6PG – 6-phosphogluconate, Ru5P – ribulose-5-phosphate, R5P – ribose-5-phosphate, X5P-xylose-5-phosphate, S7P – sedoheptulose-7-phosphate, DHAP – dihydroxyacetone phosphate, E4P – erythrose-4-phosphate, F6P – fructose-6-phosphate, GAP – glyceraldehyde-3-phosphate, THF – tetrahydrofolate, TCA - tricarboxylic acid cycle.

Here, we demonstrate that disrupting the canonical PPP in a double *zwf talAB* mutant, leading to aPPP, heightens the lethal action of diverse bactericidal antibiotics on *E. coli*. We propose that the distinctive ribose-5-phosphate metabolism in the *zwf talAB* mutant underlies the increased killing effect of antibiotics. This suggestion is supported by identification of mutations and factors mitigating the hyper-susceptibility of the *zwf talAB* mutant.

## RESULTS

### Disruption of the canonical PPP increases the killing efficiency of various antibiotics

Inactivating the oxidative branch of the PPP by deleting the *zwf* gene, which encodes glucose 6-phosphate dehydrogenase, did not impede *E. coli* growth. This is because pentose phosphates could still be synthesized from fructose-6-phosphate and glyceraldehyde-3-phosphate through the non-oxidative branch of PPP, utilizing transketolases and transaldolases (16). Cells deficient in glucose 6-phosphate dehydrogenase, along with inactivated transaldolases *(ΔtalAB*), remained viable due to compensatory activities of aldolase and fructose-1,6-phosphatase acting as sedoheptulose-1,7-phosphatase encoded by *glpX* gene (16, 17). Indeed, we demonstrated that the double *Δzwf ΔtalAB* mutant exhibits an increased level of aldolase activity (*SI Appendix,* Fig. S1A) and high level of *glpX* gene expression (*SI Appendix,* Fig. S1B). Accordingly, inactivation of *glpX* in the background of *Δzwf ΔtalAB* leads to inability to form colonies (*SI Appendix,* Fig. S2). Consequently, oxidative branch PPP-deficient cells spontaneously transitioned to anabolic PPP (aPPP) to generate pentose phosphates via the intermediate metabolite sedoheptulose-1,7-phosphate (Fig. 1B).

Remarkably, aPPP cells *(Δzwf ΔtalAB*) exhibited drastically diminished survival after treatment by various antibiotics, including quinolones, beta-lactams, and aminoglycosides, as measured by counting colony forming units (CFUs) using drug-free medium (Fig. 2A), with little effect on the minimal inhibitory concentration (MIC) (Table S1). Notably, this hypersensitivity was observed only when both branches of the PPP were inactivated, i.e., when Δ*zwf* and Δ*talAB* were combined (Fig. 2A and B). Furthermore, strains deficient in *zwf* and *talAB* displayed hyper-susceptibility to exogenous oxidants such as hydrogen peroxide and paraquat compared to WT cells (Fig. 3A and B). Notably, both Δ*zwf* and *Δzwf ΔtalAB* mutants exhibited approximately the same sensitivity to paraquat, while in the *Δzwf ΔtalAB* mutant we observed about 1-log enhancement in killing by H_2_O_2_ compared to the Δ*zwf* mutant (Fig.3 A and B). Also, Δ*zwf* and the double mutant *Δzwf ΔtalAB* showed a significant increase in spontaneous cell death in the exponentially growing culture (Fig. 3C). However, no differences were found in cell death in overnight cultures of the mutants (*SI Appendix,* Fig. S3).

**Fig. 2.**
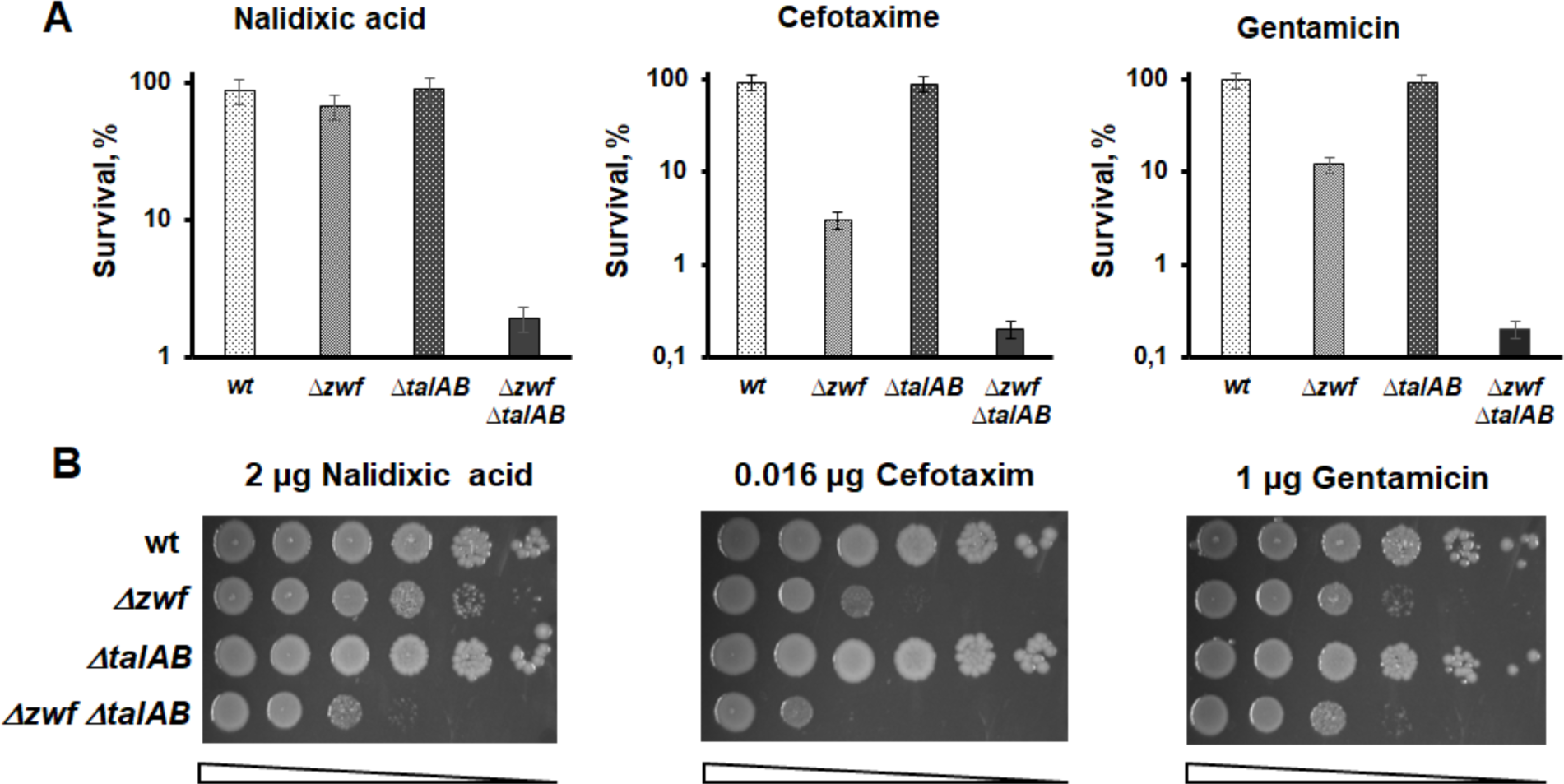
Sensitivity of *E. coli zwf talAB* mutants to different classes of antibiotics. (**A**) Overnight cultures of indicated *E. coli* strains were diluted with fresh LB 1:100 and grown to ∼2 × 10^7^. Suspensions of *E. coli* cells treated with the antibiotics nalidixic acid, gentamicin, and cefotaxime, at 5× MIC for 1 hour. Cell survival was determined by counting cfu and is shown as the mean ± SD from three independent experiments. (**B**) Representative efficiencies of colony formation of WT (MG1655) and mutant E. coli cells in the presence of various antibiotics (2 µg nalidixic acid, 1 µg gentamicin, 16 ng cefotaxime). Cells were spotted on LB agar plates in serial 10-fold dilutions and incubated at 37 °C for 24 h.

**Fig. 3.**
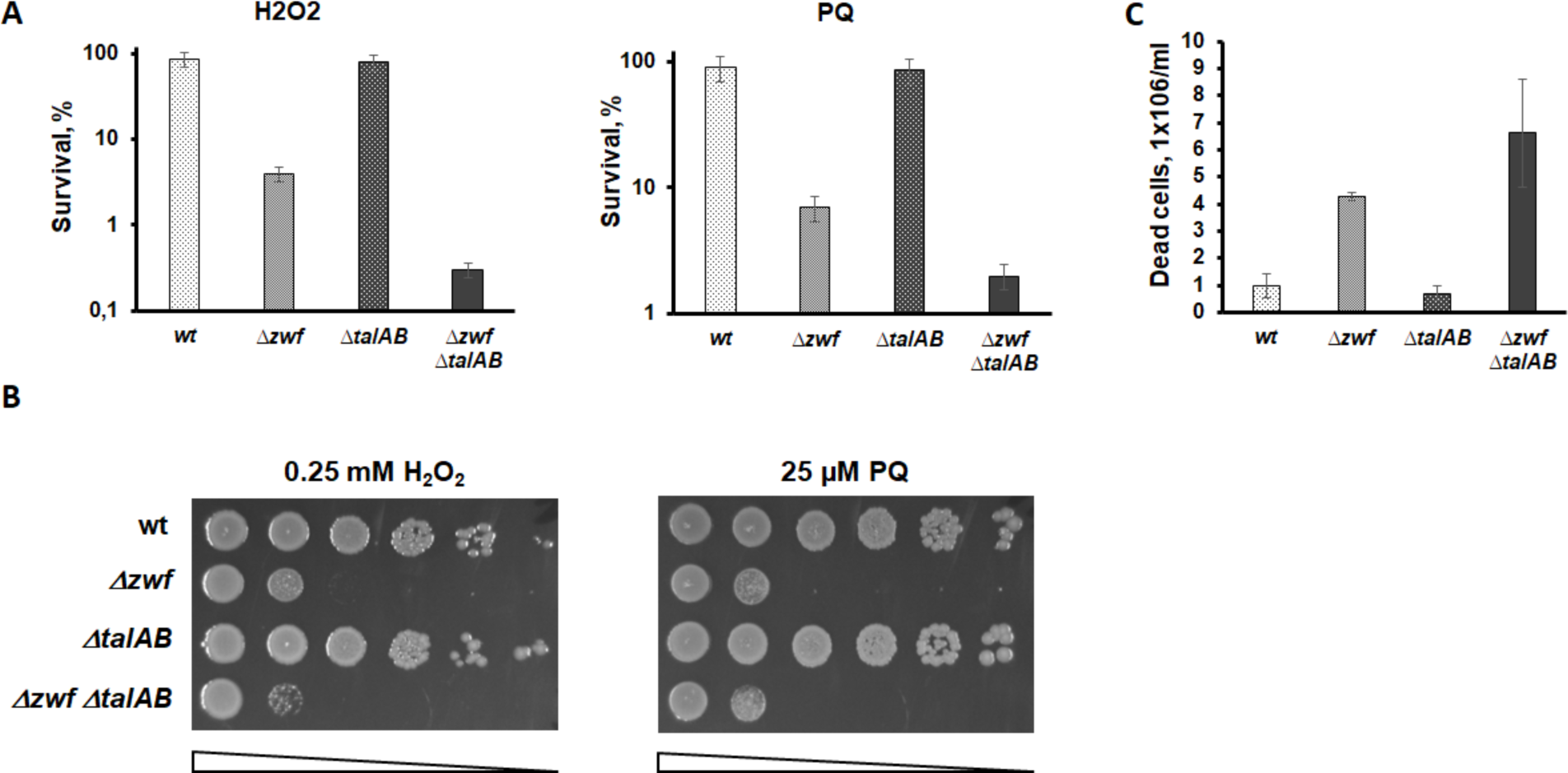
Deletion of *zwf talAB* sensitizes cells to oxidative stress. (**A**) Overnight cultures of indicated *E. coli* strains were diluted with fresh LB 1:100 and grown to ∼2 × 10^7^. H_2_O_2_ was added to 1.5 mM or 100 µM paraquat for 10 min. Cell survival was determined by counting cfu and is shown as the mean ± SD from three independent experiments. (**B**) Representative efficiencies of colony formation of WT (MG1655) and mutant *E. coli* cells in the presence of 0.25 mM H_2_O_2_ or 25 µM paraquat. Cells were spotted on LB agar plates in serial 10-fold dilutions and incubated at 30 °C for 24 h. (**C**) The number of dead cells in the cell population was determined by flow cytometry using propidium iodide. Cells were grown in an LB medium to an optical density of 0.4 and then washed twice with phosphate-buffered saline (1xPBS), centrifuged, the supernatant was removed, and cells were resuspended in 100 μL of PBS. Propidium iodide was added at a concentration of 10 μg/mL per minute before the start of the analysis.

### Disruption of the canonical PPP leads to cellular redox imbalance

Oxidative stress has been implicated in mediating antibiotic toxicity (3, 18, 19). Therefore, to elucidate the hypersensitivity of aPPP cells to antibiotic, we investigated the redox status of the separately mutations *Δzwf*, *ΔtalAB* and double mutant *Δzwf ΔtalAB*. We observed a significant accumulation of endogenous reactive oxygen species (ROS) in in *Δzwf* and *Δzwf ΔtalAB* mutants, as evidenced by the ROS-specific dye dihydrorhodamine 123 (Fig. 4A). This finding is further supported by the activation of the oxidative stress-inducible *soxS* promoter (Fig. 4B). Furthermore, aPPP cells exhibited reduced levels of the major endogenous antioxidant glutathione (Fig. 4C). Another notable redox-related feature of aPPP cells is elevated levels of NADPH (Fig. 4D), indicating a global redox imbalance upon inactivation of the oxidative branch in *Δzwf* and *Δzwf ΔtalAB* cells.

**Fig. 4.**
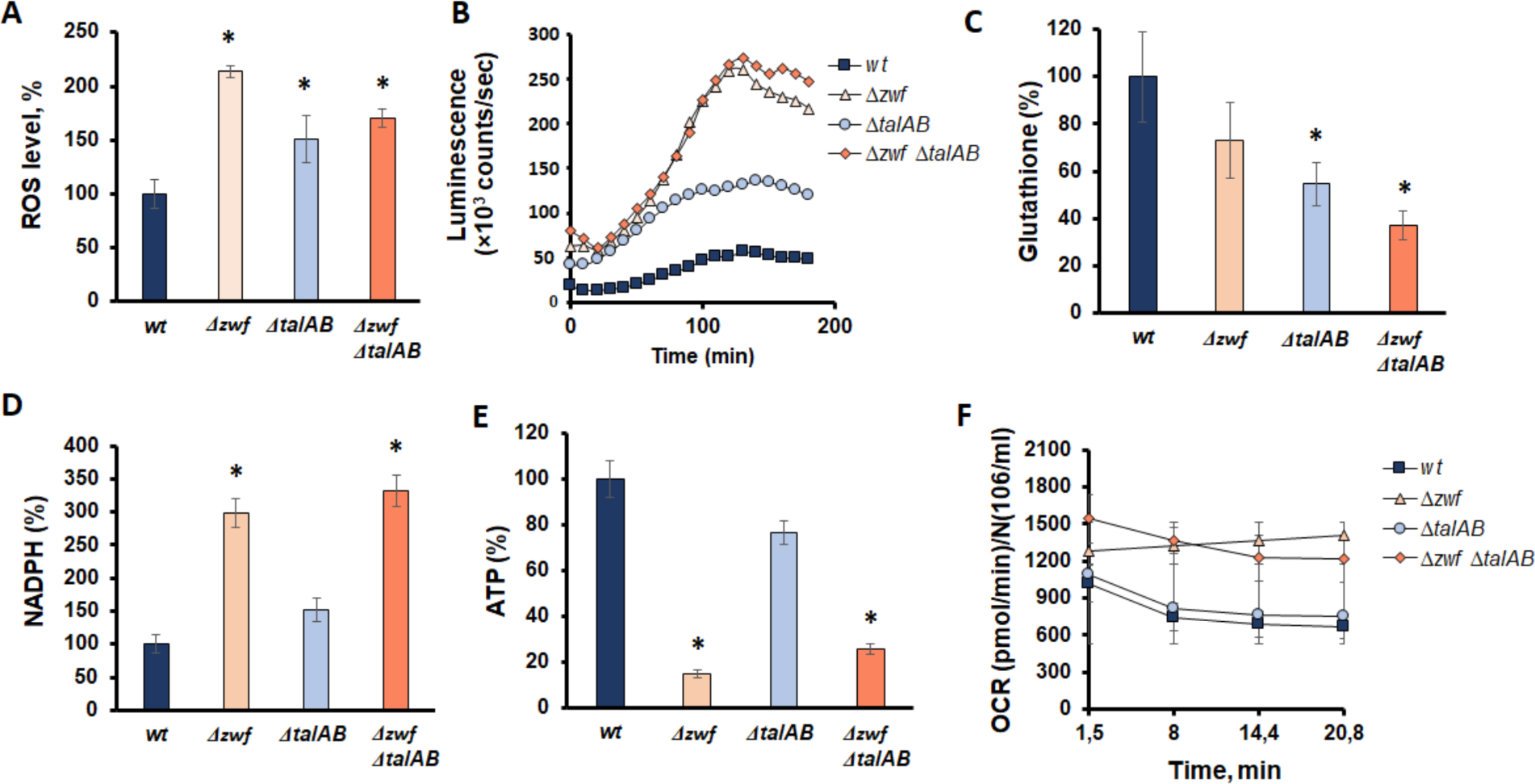
Phenotypic characteristics of Δ*zwf* Δ*talAB* mutants: activity of the soxS promoter (**A**), ROS formation (**B**), content of NADPH (**C**), glutathione (GSH) (**D**), ATP (**E**), and basal oxygen consumption rate (**F**). Mean values ± SD from at least three independent experiments are shown. * – p < 0.05, compared to the wild-type cells.

We hypothesized that in the absence of PPP, a compensatory NADPH-generating energy-consuming process could sustain high levels of NADPH, potentially explaining the observed drop in ATP content (Fig. 4E). Indeed, the hyperactivity of the serine-glycine pathway associated with purine biosynthesis may account for the increased NADPH levels in G6PDH null cells, as demonstrated by their return to normal levels upon inactivation of serine hydroxymethyltransferase (*glyA*) in Δ*zwf* cells (*SI Appendix,* Fig. S1C). Importantly, deletion of *glyA* in the wild-type parent strain did not significantly alter NADPH levels. It should be noted that all observed redox perturbation seems to be caused by the inactivation of *zwf* gene, while the deletion of *talAB* genes has no effect on redox parameters except for a decrease in intracellular glutathione content (Fig.4C).

### Ribose-5-phosphate overflow underlies antibiotics hypersensitivity

Under antibiotic exposure, *Δzwf ΔtalAB* cells are expected to accumulate ribose-5-phosphate due to the unidirectional nature of the anabolic pentose phosphate pathway (aPPP) process (Fig. 1). We hypothesize that the hypersensitivity of aPPP cells to antibiotics and the observed changes in redox status may arise from an imbalance between ribose-5-phosphate synthesis and its metabolic consumption. Processes that increase anabolic activity involving ribose-5-phosphate, as well as those that decrease ribose-5-phosphate biosynthesis, are anticipated to alleviate the *Δzwf ΔtalAB* phenotypes.

A primary metabolic process utilizing ribose-5-phosphate is nucleotide biosynthesis. Remarkably, derepression of the purine biosynthetic regulon via PurR repressor inactivation enhances the tolerance of *Δzwf ΔtalAB* cells to antibiotics (Fig. 5A) and oxidants (Fig. 5B).

**Fig. 5.**
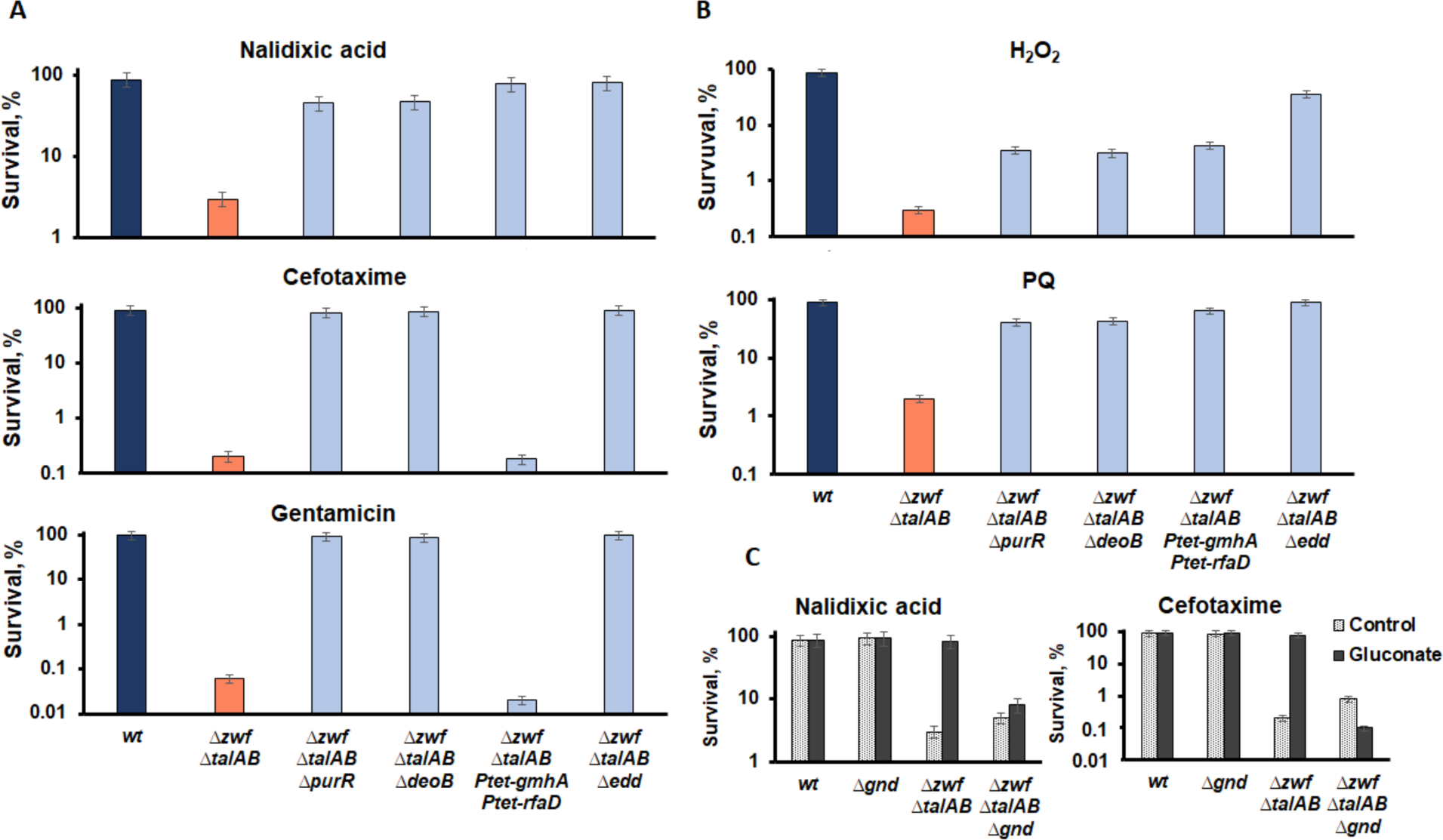
Suppression of aPPP phenotypes. Hypersensitivity to antibiotics (**A**) and redox imbalance (**B**) can be suppressed by stimulating ribose-5-phosphate utilization via enhanced purine metabolism (Δ*purR*) or ribose-5-phosphate incorporation into the cell wall (Ptet-*gmhA* Ptet-*rfaD*), or by limiting the formation of ribose-5-phosphate from nucleosides (*deoB*). Reactivation of PPP via blocking the Entner-Doudoroff pathway (Δ*edd*) completely suppresses the sensitivity of the *zwf talAB* mutant. (**C**) Addition of sodium gluconate to the growth medium restores antibiotic tolerance of the *Δzwf ΔtalAB* mutant to the level of wild-type cells. Deletion of the *gnd* gene removes the suppressive effect of sodium gluconate. Overnight cultures of indicated *E. coli* strains were diluted with fresh LB 1:100 and grown to ∼2 × 10^7. Suspensions of *E. coli* cells treated with the antibiotics at 5 × MIC or H_2_O_2_ or paraquat for 1 hour. Cell survival was determined by counting cfu and is shown as the mean ± SD from three independent experiments.

Furthermore, activation of cell wall biosynthesis, which consumes a significant portion of cellular pentose phosphates, also mitigates the hypersensitivity of *Δzwf ΔtalAB* to antibiotics and oxidants (Fig.5 A and B and *SI Appendix,* Fig. S4). The initial step in ADP-heptose formation for lipopolysaccharide (LPS) envelope biogenesis is catalyzed by D-sedoheptulose 7-phosphate isomerase (*gmhA*), followed by subsequent steps involving ADP-L-glycero-D-mannoheptose 6-epimerase (*rfaD*) (20). Overexpression of *gmhA* and *rfaD* genes increases ribose-5-phosphate flux into sedoheptulose and subsequently into cell wall LPS (21). Consequently, placing chromosomal copies of *gmhA* and *rfaD* under the control of the strong constitutive P_tet_ promoter effectively suppresses Δ*zwf* Δ*talAB* cells’ hypersensitivity to nalidixic acid and oxidants (Fig. 5A and B and *SI Appendix,* Fig. S4). However, the opposite effect is observed with respect to cefotaxime and gentamicin (Fig. 5A), suggesting interference with these antibiotics’ actions by overexpressed genes involved in cell envelope biosynthesis.

In rich LB medium, the *deo* operon, responsible for purine and pyrimidine nucleoside catabolism, significantly contributes to the ribose-5-phosphate pool (17, 22). Thus, reducing nucleoside catabolism was expected to alleviate aPPP cells’ sensitivity to antibiotics. The final step in nucleoside degradation is the conversion of ribose-1-phosphate to ribose-5-phosphate, catalyzed by phosphopentomutase (*deoB*). As anticipated, *deoB* gene deletion strongly mitigates *Δzwf ΔtalAB* cells’ hypersensitivity to oxidative stress and antibiotics (Fig. 5A and B and *SI Appendix,* Fig. S4).

Therefore, strategies aimed at to ribose-5-phosphate accumulation, such as inhibiting its utilization or promoting its formation from nucleosides, hold promise for enhancing redox imbalance and reducing antibiotic tolerance.

### Deactivation of aPPP by gluconate restores antibiotics tolerance

The deletion of the *zwf* gene prevents ribose synthesis from glucose-6-phosphate via the oxidative branch of the pentose phosphate pathway (PPP). However, the downstream part of this pathway, involving the decarboxylation of phosphogluconate to form ribulose-5-phosphate and reduced NADP^+^, remains intact. Therefore, we hypothesized that excessive exogenous gluconate could “reactivate” the PPP in *Δzwf ΔtalAB* cells by promoting its final step. Indeed, the addition of sodium gluconate to the growth medium restored antibiotic tolerance of the *Δzwf ΔtalAB* mutant to wild-type levels (Fig. 5C). This suppressive effect of sodium gluconate was completely abolished by the deletion of the *gnd* gene encoding phosphogluconate dehydrogenase, the enzyme responsible for the final step of oxidative branch of PPP (Fig. 5C).

Excess phosphogluconate in the cell can also be achieved by blocking the Entner-Doudoroff pathway (23), where the initial step is catalyzed by phosphogluconate dehydratase 1(*edd*). Inactivation of the *edd* gene in the *ΔtalAB Δzwf* strain completely suppressed the sensitivity of *ΔtalAB Δzwf* cells to all antibiotics used in our study (Fig. 5A). These findings suggest that gluconate metabolism can deactivate the anabolic PPP in PPP-deficient *Δzwf ΔtalAB* cells, effectively mimicking normal PPP functioning.

It is noteworthy that the suppression of hypersensitivity of *Δzwf ΔtalAB* cells to antibiotics through *purR*, *deoB*, Ptet-*lpcA* Ptet-*rfaD*, and *edd* mutations consistently accompanied the restoration of intracellular glutathione content (*SI Appendix,* Fig. S5A) and ATP levels (*SI Appendix,* Fig. S5C), nearly to the level of wild-type cells. Moreover, the suppression of antibiotic hypersensitivity was associated with a reduction in the level of reduced NADPH (*SI Appendix,* Fig. S5B), suggesting an inverse correlation between intracellular glutathione and NADPH levels.

## DISCUSSION

It has been observed that when both branches of the canonical pentose phosphate pathway (PPP) are shut down, microorganisms can switch to alternative anabolic synthesis of pentose phosphates. Three glycolytic enzymes—sedoheptulose-7-phosphate phosphokinase (Pfk), sedoheptulose-1,7-bisphosphate phosphatase (GlpX), and aldolase A (FbaA) are required for the construction of a new Tal-less non-oxidative branch of the PPP (Fig. 1) (16, 17). In the absence of the oxidative branch due to deletion of the key *zwf* gene, the inverted Tal-less non-oxidative branch becomes the sole pathway for synthesizing pentose phosphates (Fig. 1). It is important to note that in the anabolic PPP (aPPP), the synthesis of pentose phosphates is unidirectional, as ATP is utilized to phosphorylate sedoheptulose-7-phosphate. Our findings indicate that the activation of aPPP in *E. coli* leads to enhanced sensitivity to oxidative stress and several of classes of bactericidal antibiotics. Furthermore, aPPP causes significant redox imbalance, characterized by elevated endogenous ROS and NADPH levels, and decreased levels of glutathione and ATP (Fig. 4). The activation of aPPP also leads to increased respiration, indicating intensified metabolic activity (Fig. 4F). The deletion of *zwf*, which is part of the redox-sensing SoxRS regulon, appears to decouple the oxidative stress response from the coordinated generation of reducing equivalents (NADPH) and pentose phosphates biosynthesis. A specific feature of aPPP is the futile cycling of sedoheptulose-1,7-bisphosphate and sedoheptulose-7-phosphate, which consumes ATP in parallel with pentose phosphates utilization in glycolysis (Fig. 1).

Recent studies have highlighted the important regulatory role of ribose-5-phosphate plays in coordinating nucleotide recycling and amino acid synthesis in the stringent response (24–26). Furthermore, in our previous study, we found that impaired efflux of the sedoheptulose-7-phosphate (via *gmhA* deletion) from the PPP leads to a significant increase in bacteria sensitivity to antibiotics, which can be suppressed by activating purine biosynthesis through *purR* deletion (27). Further experiments aimed to elucidate the connections between pentose phosphate content and antibiotic hypersensitivity of the mutant. The action of any antibiotic inhibits all anabolic biosynthetic processes associated with pentose phosphates consumption, thereby inducing coordination of synthesis and reutilization of pentoses through cyclization of the interconversion of the sedoheptulose-1,7-bisphosphate and sedoheptulose-7-phosphate pair. Strengthening the processes of pentose phosphate consumption by promoting purine biosynthesis via *purR*, or stimulating cell envelope synthesis through Ptet-*gmhA* and Ptet-*rfaD*, or limiting ribose-5-phosphate formation as a result of nucleoside degradation through *deoB*, leads to the restoration of antibiotic tolerance and specific physiological parameters in aPPP cells (Fig. 5, Fig. S2 and S3).

Of particular interest is the suppression of sensitivity by increasing the intracellular concentration of phosphogluconate, as it is directly based on the mechanism of metabolic futile cycling. It is known that the rate of ATP-dependent reaction of the futile cycle slightly increases, while the rate of the reversed reaction changes significantly; thus, the final rate of the direct reaction will be significantly increased. The futile cycling acts as an enhancer of slight changes in the rate of the direct reaction (28). Inactivation of the Entner-Doudoroff pathway (*Δedd*) or shifting the metabolic equilibrium by the addition of exogenous phosphogluconic acid promotes the flux of pentose phosphates to glycolysis and thus represses the alternative process of pentose phosphates synthesis from glycolysis, preventing the occurrence of the futile cycle in *Δzwf ΔtalAB* cells (Fig. 4C).

In summary, the findings presented in this study suggest that the induction of unidirectional aPPP is a critical factor in the ability of bacteria to tolerate antibiotics. Therefore, manipulating the two ways of pentose phosphates generation (oxidative branch of canonical PPP and aPPP pathway) may have therapeutic potential to enhance antibiotic efficacy and combat antibiotic resistance.

## Materials and Methods

### Strains and Growth Conditions

All *E. coli* strains utilized in this work are listed in *SI Appendix,* Table S2. BW25113 and its derivatives (single gene deletion mutants) were obtained from the E. coli Keio Knockout Collection (Thermo Scientific) (29). P1 transduction was employed to introduce mutations into new strains (30). When necessary, Cam or Kan drug resistance markers were excised from strains using the FLP activity of pCP20, followed by loss of the plasmid at the nonpermissive temperature (31). All mutations were verified by PCR and gel analysis. DNA manipulation and the transformation of *E. coli* strains were performed according to standard methods (32). LB complete medium was used for the routine growth of E. coli. When appropriate, antibiotics were added at 40 μg/mL (for Km), 30 μg/mL (for chloramphenicol), and 100 μg/mL (for ampicillin). For solid medium, 1.5% agar was added.

### Ptet-gmhA and Ptet-rfaD Constructions

To construct the *gmhA* and *rfaD* overexpression strains, the native promoter of these genes was substituted by PLtet-O1 (33). Briefly, the P Ltet-O1-attL-CmR-attR cassette integrated into the AM3009 strain *mstA* gene (34) was amplified with primers 5′-gcacttcaggtcaaaaagtcctggtcatagcacctgcgctcaagttagtataaaaaagct-3′ and 5′-ttcgttcag-ttcgttacgaataagatcctggtacatggtacctttctcctctttaatga-3′ (for *gmhA* gene); and 5′-ttcacatgcaaaaccaacatccgccatgaaggactacgctcaagttagtataaaaaagct-3 and 5′-gccgat-aaagcccgcgccgccggtaacgatgatcatggtacctttctcctctttaatga-3′ (for *rfaD* gene). The first set of primers contained the upstream region of *gmhA* and *rfaD* genes and the sequence of attR, while the second set of primers contained the coding region of corresponding genes and the sequence of PLtet-O1. The PCR fragments were transformed into MG1655 containing pKD46 (31). CmR clones were tested in the presence of the PLtet-O1-attL-CmR-attR cassette by PCR with primers 5′-aacaaagctcacattgttgct-3′ and 5′-gcgctgaatggcgtgaatatt-3′ (for *gmhA* gene); and 5′-atcggaatattgatactaaagc-3′ and 5′-aatatcggtgatgcctttatc-3′ (for *rfaD* gene). All constructs were sequenced for verification and introduced into corresponding chromosomal loci according to ref. 35. All strains bearing Ptet constructs do not contain the *tetR* gene and, therefore, exhibit constitutive expression of target genes.

### Determination of Sensitivity to Antibiotics

Overnight bacterial cultures were diluted 100 times with fresh LB medium and grown to OD_600_≈0.5–0.6 under aerobic conditions at 37°C. Cell suspensions were aligned in optical density, and a series of tenfold dilutions were prepared in a 96-well plate. Ready dilutions were plated on plates with a rich agar medium containing various concentrations of the studied antibiotics: nalidixic acid, moxifloxacin, cefotaxime, gentamicin, and rifampicin. The dishes were incubated overnight at 37°C, and the result was photographed on a GelCamera M-26XV detection system. For antibiotic survival assays, overnight bacterial cultures were diluted 100-fold and grown at 37°C to ∼2 × 10^7^ cells per mL, treated with the indicated concentration of nalidixic acid, cefotaxime, or gentamicin, and after 90 minutes of incubation, samples were diluted and plated on LB agar and incubated at 37°C for 24 hours. Cell survival was determined by counting cfu and is shown as the mean ± SD from three independent experiments.

### Detection of pSoxS Activity

The pSoxS::lux plasmid (lux biosensor) containing the *Photorhabdus luminescens* lux operon genes under the control of the *soxS* gene promoter was used as a reporter system for detecting the formation of reactive oxygen species (35). All studied strains were transformed with this plasmid. Overnight cultures of the strains were diluted 100 times and grown to OD_600_≈0.2. Suspensions were normalized by optical density and transferred to a 96-well plate in 200 μL. Detection of bioluminescence was carried out in a tablet reader Tecan Spark at a wavelength of 490 nm for 3 hours at room temperature.

### Determination of Viability of Cells and ROS Using Flow Cytometry

Cells were grown in complete medium to an optical density of 0.4 and then washed twice with phosphate-buffered saline (1xPBS), centrifuged, the supernatant was removed, and cells were resuspended in 100 μL of PBS. Cell parameters were analyzed using flow cytometry on a BD LCR Fortessa flow cytometer (Becton Dickinson, Franklin Lakes, New Jersey, USA). The cell population for analysis was selected according to the parameters of forward (FSC) and side scattering (SSC), which characterize the size and granularity of cells. The percentage of dead cells in the cell population was assessed using propidium iodide (Ex/Em = 535/617 nm, Sigma-Aldrich, St. Louis, Missouri, USA), which was added to the cells at a concentration of 10 μg/mL per minute before the start of the analysis. Propidium iodide penetrates the cells with a damaged membrane and, after binding to DNA, has a bright fluorescence in the red region of the spectrum. The ROS level was assessed using the dye Dihydrorhodamine 123 (DHR123) (Ex/Em = 507/525 nm, ThermoFisher Scientific, Waltham, Massachusetts, USA) (36), which was added to the cells to a final concentration of 7.5 µM, and the cells were incubated for an hour in the dark at 37°C.

### Measurement of NADPH Level

Measurement of NADPH levels was performed using a fluorimetric NADP/NADPH Assay Kit (Abcam). Cells were grown to OD600≈0.5 in a thermostated shaker at 37°C. Preparation of cell extracts, as well as all subsequent manipulations, was carried out according to the manufacturer’s instructions attached to the kit. Sample fluorescence was detected in a Tecan Spark plate reader at Ex/Em = 540/590 nm. The results obtained were related to the optical density of the OD600 culture.

### Measurement of Intracellular Glutathione Levels

Quantitative determination of the level of intracellular glutathione was carried out by the modified Tietz method (37). 100 µl of overnight culture was transferred to 10 ml of fresh LB medium and grown to OD_600_≈0.5. Cells from a 5 ml suspension were pelleted by centrifugation and suspended in a lysis buffer: 0.1M potassium phosphate buffer with 5 mM EDTA disodium salt, pH 7.5, supplemented with 0.1% Triton X100. The cells were homogenized with a pestle and centrifuged at 11,000 rpm for 5 minutes at 4°C. Supernatant was transferred into clean test tubes. The reaction mixture of 60 µl of cell extract, 120 µl of DTNB (0.3 mg/ml) and GOR (1.5 units/ml) and 60 µl of NADPH (0.6 mg/ml) was incubated for 3 minutes at room temperature, after which adsorption was measured at 412 nm. GSH (Sigma Aldrich, USA) was used to construct a calibration curve. The obtained values were referred to the optical density of the cultures.

### ATP Level Measurement

The intracellular ATP level was determined by the luminescence method using the Adenosine 5’-triphosphate (ATP) Bioluminescent Assay Kit (Sigma Aldrich, USA). 5 ml of cell suspension grown to OD600≈0.5 was washed twice with saline and suspended in 2 ml of saline. Preparation of the reaction mixture and measurement of luminescence was carried out as described in the manufacturer’s instructions. The obtained values were related to the optical density of the culture and expressed as a percentage. The level of ATP in the wild-type strain was taken as 100%.

### Bacterial Respiration

Seahorse XFe24 Analyzer was used to quantitate oxygen consumption rates (OCRs) (15). An overnight of *E. coli* cells was diluted 1:100 into fresh LB media and grown to an OD600 of ≈0.4. Cells were diluted to 5× the final OD, and 500 μL of diluted cells was added to XF Cell Culture Microplates precoated with poly-D-lysine (PDL) (5). Cells were centrifuged for 10 min at 1,400 × g to attach them to the precoated plates. Basal OCR was measured for four cycles (20 min). OCR was normalized to the number of viable cells quantitated using flow cytometry on a BD LCR Fortessa flow cytometer (Becton Dickinson, Franklin Lakes, New Jersey, USA).

### RNA Extraction and qRT-PCR

E. coli K-12 MG1655, TS011, TS013, and TS015 cells were grown until OD_600_ ≈0.5 and total RNA was extracted using the RNeasy Mini Kit (QIAGEN) according to the manufacturer’s protocol according to ref. 21. All RNA samples were treated with DNaseI (Fermentas). 500 ng of total RNA were reverse-transcribed with 100U of SuperScript III enzyme from the First-Strand Synthesis Kit for RT-PCR (Invitrogen) according to the manufacturer’s protocol in the presence of appropriate gene-specific primers. 1μl of the reverse transcription reactions was used as a template for real-time PCR. The gene *def* encoding peptide deformylase was used for normalization. Each real-time PCR mixture (25μl) contained 10 μl SYBR Green I PCR Master Mix (Syntol), 12 μl nuclease-free H_2_O, 1μl of 10 μM forward primer, 1 μl of 10μM reverse primer and 1 μl of cDNA template. Amplifications were carried out using DTlite S1 CyclerSystem (DNA Technology). Reaction products were analyzed using 2% agarose electrophoresis to confirm that the detected signals originated from products of expected lengths. Each qRT-PCR reaction was performed at least in triplicate and average data are reported. Error bars correspond to the standard deviation.

### Determination of Aldolase Activity

Aldolase activity was measured using the Aldolase Activity Assay Kit (Colorimetric) (Abcam). Cells were washed with cold saline and concentrated in 100 µl of Aldolase Assay Buffer to a density of 1×106 per ml. Further manipulations were performed in accordance with the manufacturer’s recommendations. The obtained values of aldolase activity were related to the concentration of total protein in the studied cell extracts.

### Statistical Analysis

The data are shown as mean ± standard deviation measures from triplicate values obtained from 3–4 independent experiments. The statistical difference between experimental groups was analyzed by one-way ANOVA with Tukey correction for multiple comparisons. Probability values (p) less than 0.05 were considered significant. Statistical analysis was performed using the GraphPad Prism 9.1.2 software (GraphPad Software Inc., San Diego, CA, USA).

## Data Availability

All study data are included in the main text and Supplemental material.

## ACKNOWLEDGMENTS.

This work was supported, in part, by Russian Science Foundation Grant 24-14-00089 (to T.S., R.S., S.S., I.P., V.M. A.M. and A.S.M.), Department of Defense Grant PR171734 (to K.S.), the Blavatnik Family Foundation, and the HHMI (to E.N.).

---

**Fig. S1.**
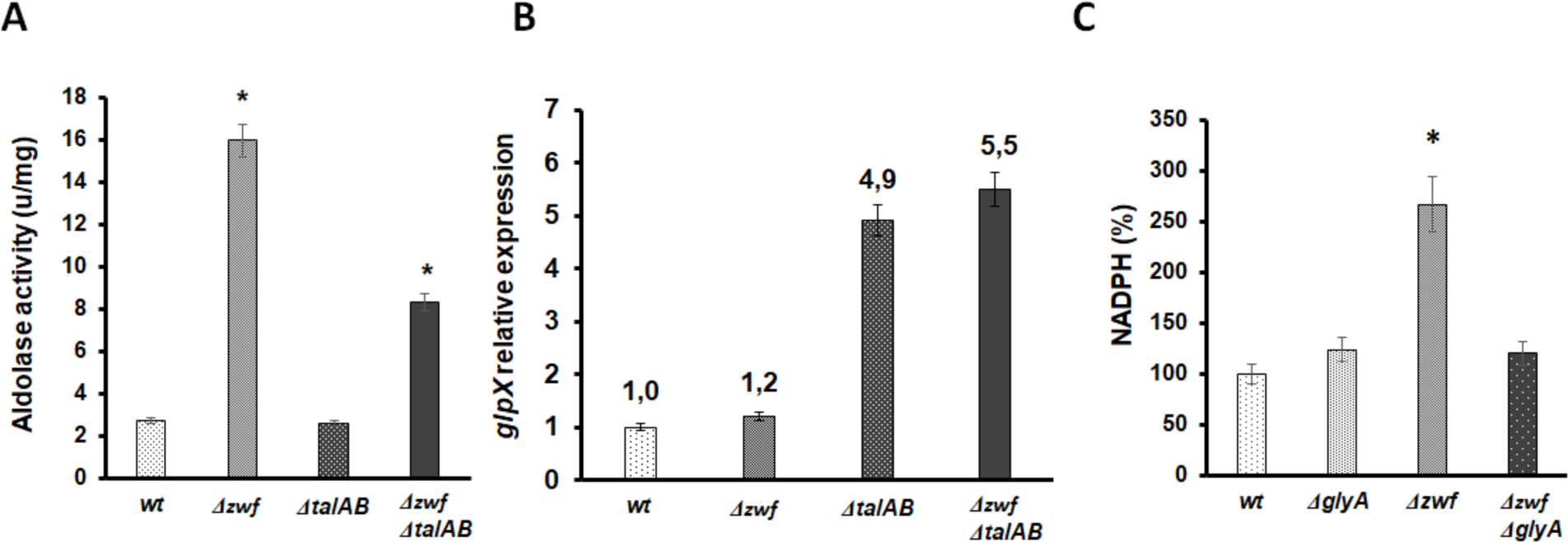
Glycolitic enzymes aldolase A, phosphatase GlpX and serine hydroxymethyltransferase are involved in aPPP. (**A**) Activity of aldolase encoded by the *fbaА* gene. Mean values ± SD from at least three independent experiments are shown. * – p < 0.05, compared to the wild-type cells. (**B**) The relative expression of the *glpX* gene in exponentially grown WT and mutant cells was measured by qRT-PCR. The relative expression (y-axis) represents the fold change of each mRNA level compared with that of the wt cells. Values are means ± SD from four experiments. (**C**) Intracellular content of NADPH in the Δ*zwf* mutant upon inactivation of the *glyA* gene. Mean values ± SD from at least three independent experiments are shown. * – p < 0.05, compared to the wild-type cells.

**Fig S2.**
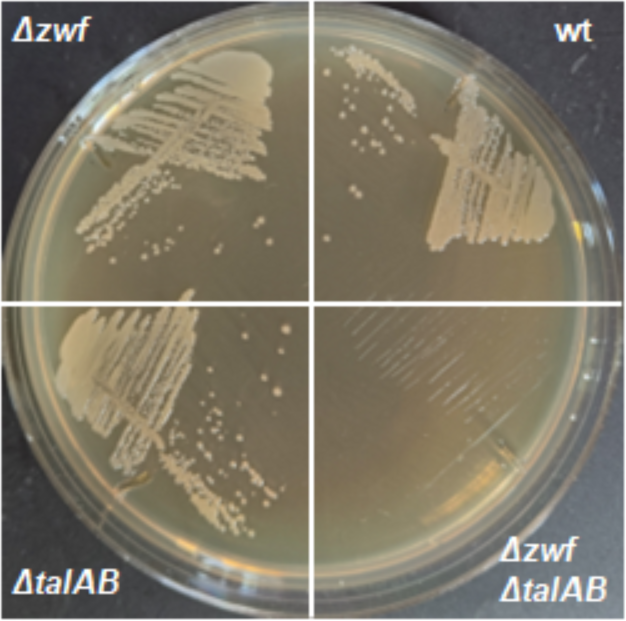
Appearance of Kan^R^ recombinants in transduction experiments. P1 grown on Keio strain JW3896 *glpX*::kan with recipient: WT, *Δzwf*, *ΔtalAB* and *Δzwf ΔtalAB*. After transduction, the cells were plated on LB medium with Kan and incubated at 37ᵒC overnight. It was not possible to obtain viable colonies for the double mutant *Δzwf ΔtalAB*. The presence of the *glpX* gene deletion was confirmed by PCR.

**Fig S3.**
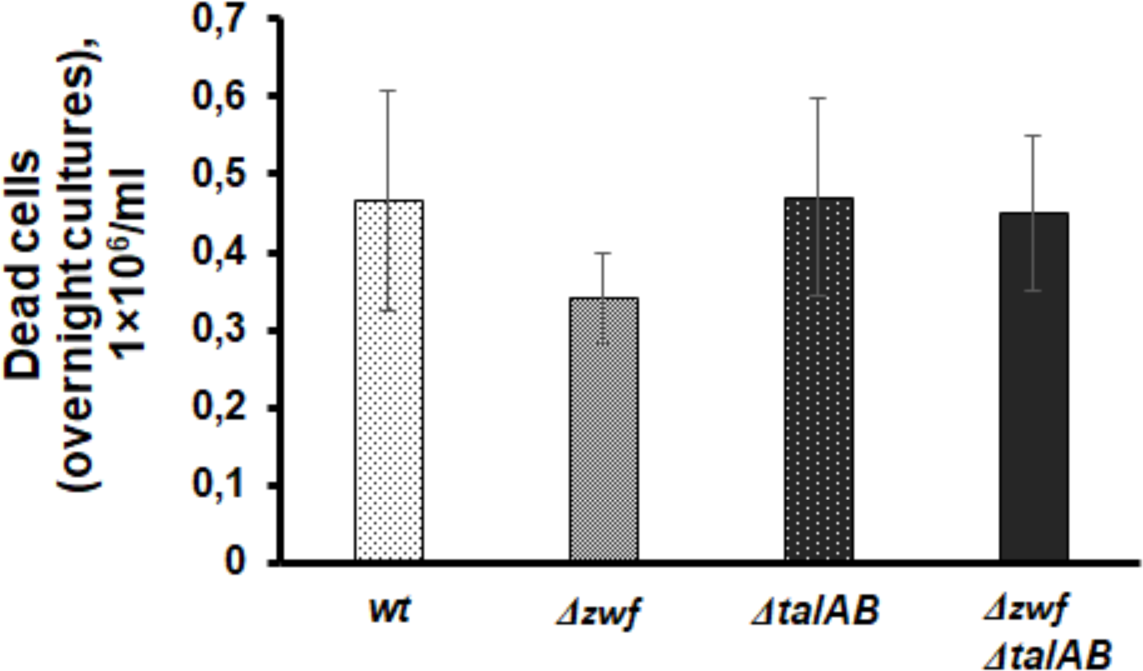
The number of dead cells in the cell population of overnight cultures was determined by flow cytometry using propidium iodide. Cells were grown in an LB medium to an optical density of 0.4 and then washed twice with phosphate-buffered saline (1xPBS), centrifuged, the supernatant was removed, and cells were resuspended in 100 μL of PBS. Cells were incubated in the presence of propidium iodide at a concentration of 10 μg/mL within 1 minute before the start of the analysis.

**Fig. S4.**
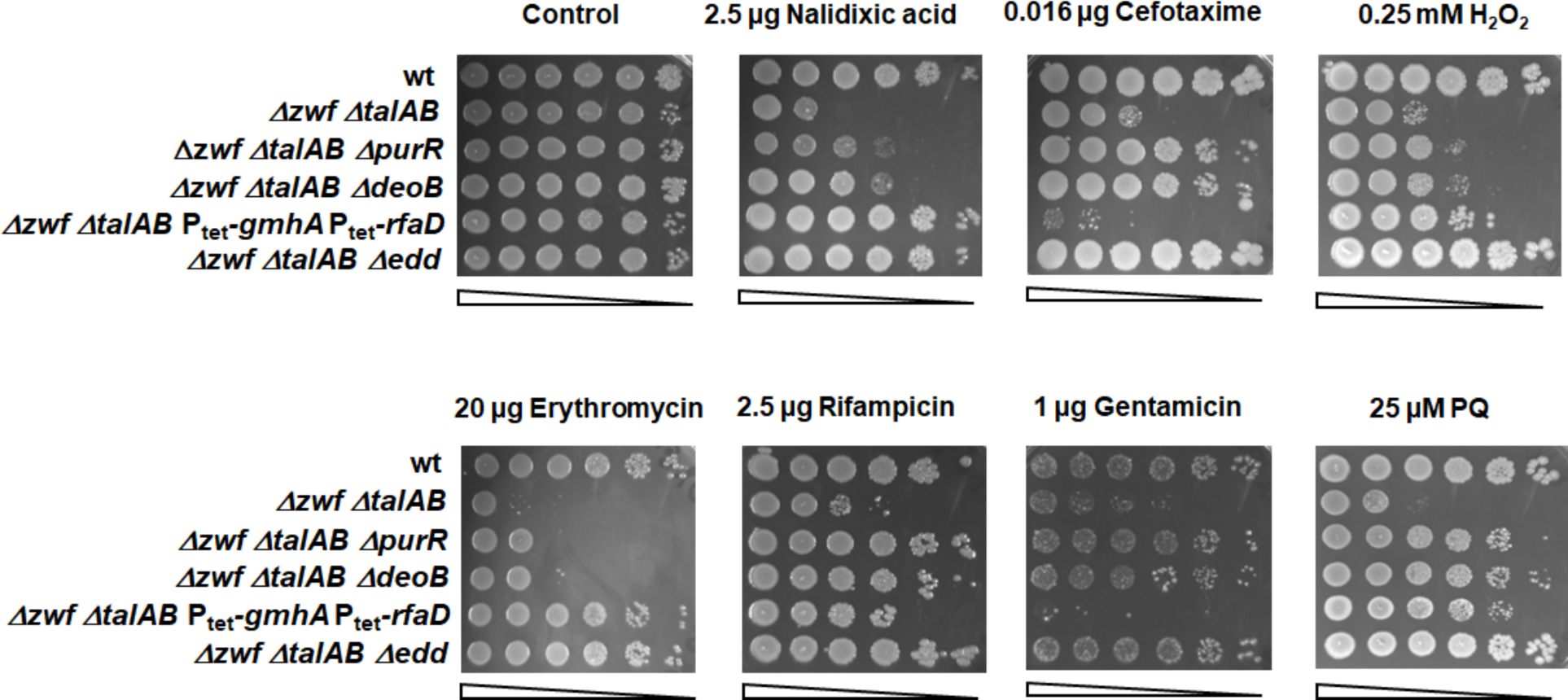
The survivability of *Δzwf ΔtalAB* cells is rescued by inactivation of *purR*, *deoB*, or *edd* gene or by *gmhA* and *rfaD* overexpression. To determine representative efficiencies of colony formation, cells were grown to OD600 ∼ 0.4, and serial 10-fold dilutions were spotted on LB agar plates containing the indicated concentrations of antibiotics or H_2_O_2_ and paraquat and incubated at 30 °C for 24 h.

**Fig. S5.**
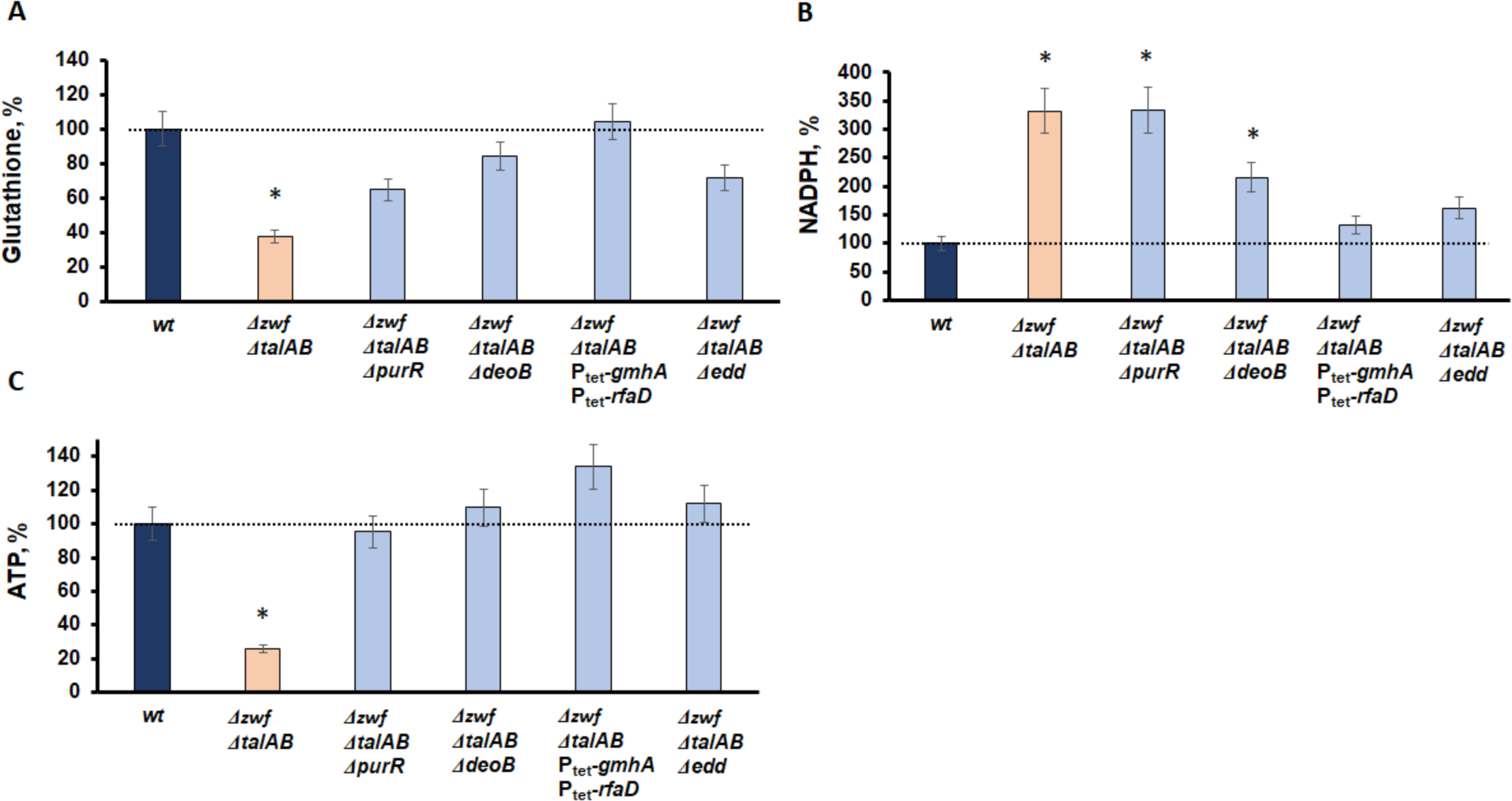
Intracellular content of glutathione (A), NADPH (B), and ATP (C) in the *Δzwf ΔtalAB* mutants against the background of *purR*, *deoB*, *edd*, and P_tet_-*gmhA* P_tet_-*rfaD* mutations. Mean values ± SD from at least three independent experiments are shown. * – p < 0.05, compared to the wild-type cells.

**Table S1.**
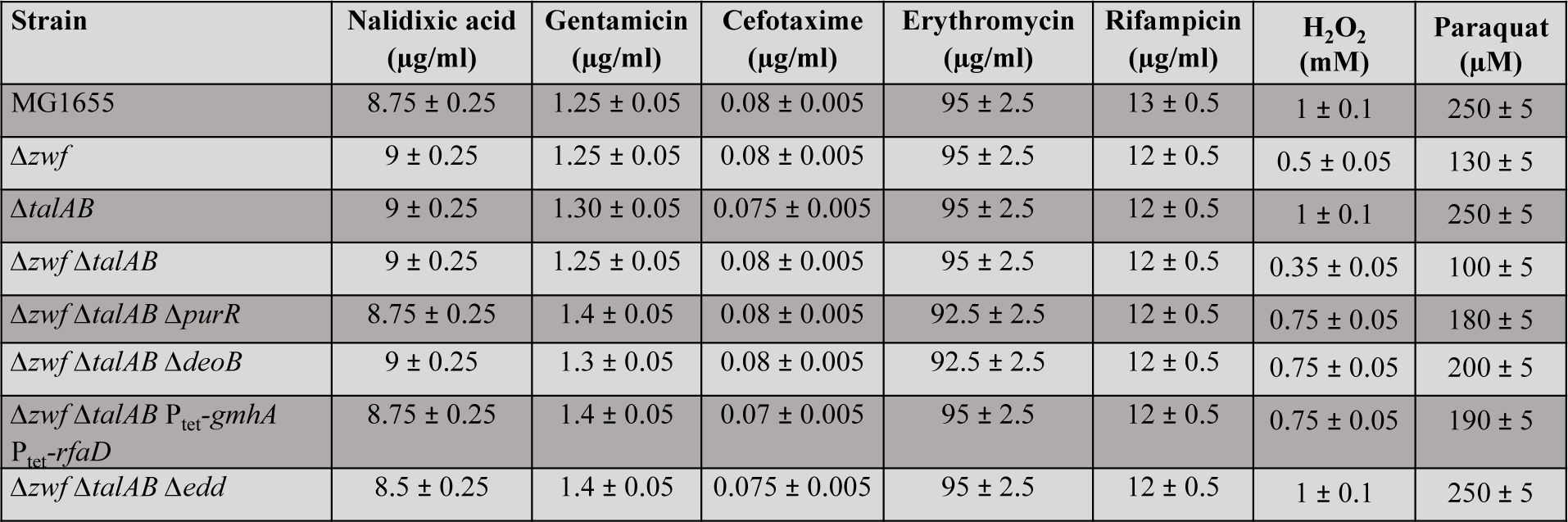
MICs for strains used in this study.

**Table S2.**
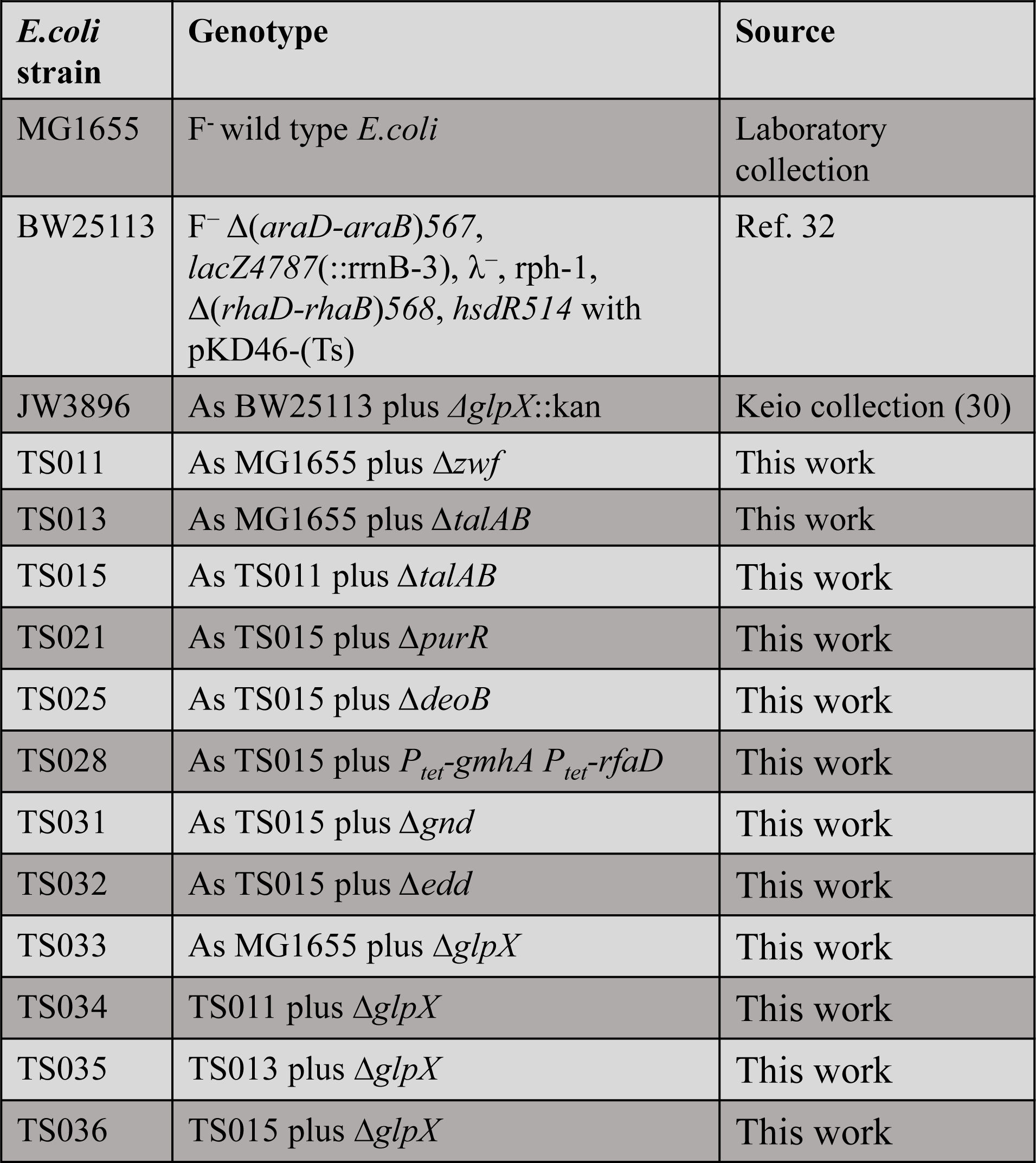
Bacterial strains.

| ***E.coli***<br>**strain** | **Genotype** | **Source** |
| --- | --- | --- |
| MG1655 | F<sup>-</sup> wild type *E.coli* | Laboratory collection |
| BW25113 | F<sup>-</sup> $\Delta(araD-araB)567$, *lacZ*4787(::rrnB-3), $\lambda^-$, rph-1, $\Delta(rhaD-rhaB)568$, *hsdR*514 with pKD46-(Ts) | Ref. 32 |
| JW3896 | As BW25113 plus $\Delta glpX::kan$ | Keio collection (30) |
| TS011 | As MG1655 plus $\Delta zwf$ | This work |
| TS013 | As MG1655 plus $\Delta talAB$ | This work |
| TS015 | As TS011 plus $\Delta talAB$ | This work |
| TS021 | As TS015 plus $\Delta purR$ | This work |
| TS025 | As TS015 plus $\Delta deoB$ | This work |
| TS028 | As TS015 plus $P_{tet}-gmhA P_{tet}-rfaD$ | This work |
| TS031 | As TS015 plus $\Delta gnd$ | This work |
| TS032 | As TS015 plus $\Delta edd$ | This work |
| TS033 | As MG1655 plus $\Delta glpX$ | This work |
| TS034 | TS011 plus $\Delta glpX$ | This work |
| TS035 | TS013 plus $\Delta glpX$ | This work |
| TS036 | TS015 plus $\Delta glpX$ | This work |

